# ALEdb 1.0: A Database of Mutations from Adaptive Laboratory Evolution Experimentation

**DOI:** 10.1101/320747

**Authors:** Patrick V. Phaneuf, Dennis Gosting, Bernhard O. Palsson, Adam M. Feist

## Abstract

Full genomic sequences are readily available, but their functional interpretation remains a fundamental challenge. Adaptive Laboratory Evolution (ALE) has emerged as an experimental approach to discover causal mutations that confer desired phenotypic functions. Thus, ALE not only represents a controllable experimental approach to systematically discover genotype-phenotype relationships, but it also allows for the revelation of the series of genetic alterations required to acquire the new phenotype. Numerous ALE studies have appeared in the literature providing a strong impetus for developing structured databases to warehouse experimental evolution information and make it retrievable for large-scale analysis. Here, the first step towards establishing this capability is presented: ALEdb (http://aledb.org). This initial release contains over 11,000 mutations that have been discovered in ALE experiments. ALEdb is the first of its kind; (1) it is a web-based platform that comprehensively reports on ALE acquired mutations and their conditions, (2) it reports key mutations using previously established trends, (3) it enables a search-driven workflow to enhance user mutation functional analysis, (4) it allows exporting of mutation query results for custom analysis, (5) it has a bibliome that describes the underlying published literature, and (6) contains experimental evolution mutations from multiple model organisms. Thus, ALEdb is an informative platform which will become increasingly revealing as the number of reported ALE experiments and identified mutations continue to expand.

## INTRODUCTION

Adaptive Laboratory Evolution (ALE) is a tool for the study of microbial adaptation. The typical execution of an ALE experiment involves cultivating a population of microorganisms in defined conditions (i.e., in a laboratory) for a period of time that enables the selection of improved phenotypes. Standard model organisms, such as *E. coli,* have proven well suited for ALE studies due to their ease of cultivation and storage, fast reproduction, well known genomes, and clear traceability of mutational events [1]. With the advent of accessible whole genome resequencing, associations can be made between selected phenotypes and genotypic mutations [2].

Beginning with a starting strain, an ALE experiment can be executed by serially passing a selected culture to a fresh flask of media (Figure 1A), enabling the strain passed to continue acquiring mutations under the experimental conditions without dilution of resources. Strains propagated during ALEs are assumed to be those that outcompeted their competition due to adaptive mutations. Additional methods to perform ALEs have been reviewed [2,3]. Whole genome comparative sequencing, or resequencing, is used to identify mutations within evolved strains relative to the evolution’s starting strain (Figure 1B). ALE experiments can additionally involve replicate ALEs: identical evolutions that are often executed in parallel. Replicate ALEs can reveal the dynamics of adaptation by enabling research into converging genotypes within an experiment [4].

**Figure 1.**
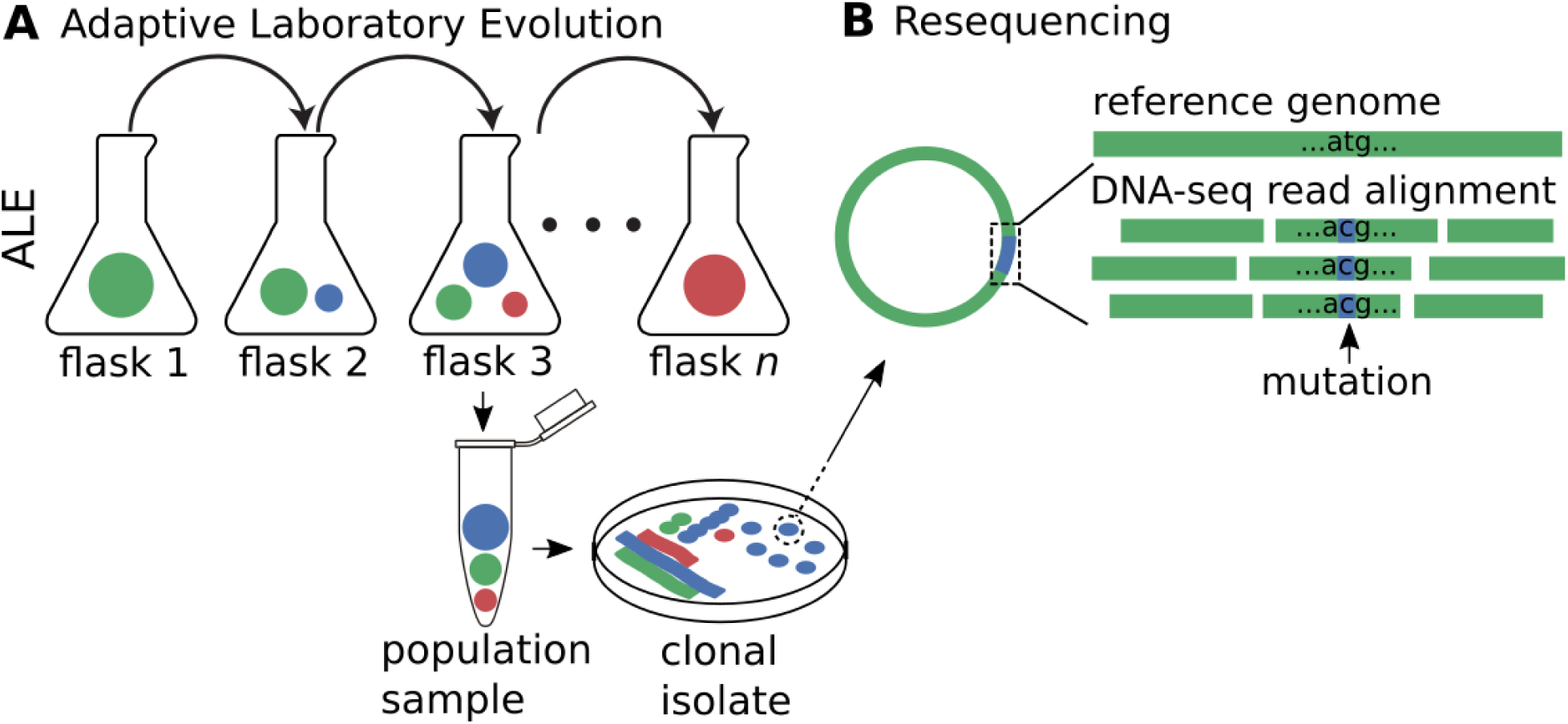
**A** An illustration of a batch ALE experiment where both a clonal and population sample are isolated from an intermediate (i.e., midpoint) flask. The petri dish represents streaking methodology for isolating a clonal colony from a population. **B** An illustration of how the resequencing process leverages a reference genome sequence and DNA-seq reads to identify mutations in an ALE sample.

ALE methods have become important scientific tools in the study of evolutionary phenomena and have contributed to research in basic discovery and applied fields. Evolutionary biologists seek to examine the dynamics and repeatability of evolution and to better understand the relationship between genotypic and phenotypic changes [5]. ALE methods, along with the plummeting cost of sequencing, has greatly enabled their efforts, resulting in a variety of insights into adaptive evolution. ALE has often demonstrated that (1) increases in fitness diminish with each new adaptive mutation [6], (2) genotypic convergence through mutations can occur on the level of functional complexes [7], and, (3) interactions between mutations may cause nonlinear fitness effects [8].

ALE methods have also been leveraged in the applied research of synthetic biology to engineer microbes for commodity, industrial, and biopharmaceutical chemical synthesis [2]. Comprehensive whole genome rational design is rarely achievable due to the complexity of biological systems [4,9]. The inability to provide for comprehensive solutions in genome engineering can result in strains which cannot maintain homeostasis, such as strains which cannot tolerate the concentrations of products they were designed to produce. ALE has been used to produce adaptive mutations that provide solutions for the gaps left by current rational genome engineering methods [10]. ALE can therefore complement rational genome engineering in the work to provide for a comprehensive whole genome solution to an application [2,9].

Accurately interpreting the results of an ALE requires the identification of causal mutations for observed adaptations. Identifying causal mutations requires a clear understanding of the mechanistic effects of mutations on cellular components and systems. Due to the complexity of cellular systems, interpreting the effects of mutations has proven to be a primary challenge in ALE [4,9]. A common approach to mutation functional analysis is a literature search on the mutation target (e.g., a given annotated ORF). Functional studies of genetic targets have traditionally served as primary resources for interpreting mutation effects, providing information on a sequence’s biological function. Published ALE results can enhance approaches to identify and understand new adaptive mutations since they describe the fitness of an allele relative to its predecessor. Researchers can therefore work to identify and understand their ALE mutations by considering published adaptive mutations in conditions similar to their own ALEs.

A review of ALE methods [2] lists 34 separate ALE studies. Each study reports on novel combinations of selection conditions and the resulting microbial adaptive strategies. Large scale analysis of ALE results data from such consolidation efforts could be a powerful tool for identifying and understanding novel adaptive mutations. No current online platform exists for ALE experimental result consolidation.

A web platform named *ALEdb* (aledb.org) has been created to meet the need for accessible consolidated ALE mutations, conditions, and publication reporting. ALEdb additionally includes features to search for specific mutations, report key mutations, and export mutation data for custom analysis. With these features, ALEdb serves to fill the gap in the field of experimental evolution for an accessible resource of consolidated experimental evolution mutations.

## RESULTS

A web platform to accelerate ALE data to knowledge

The need for consolidated and accessible ALE experiment reporting has resulted in the generation of the web platform *ALEdb* (aledb.org). Eleven published ALE experiments, with a total of four distinct strains, 532 samples and 21522 observed mutations, serve as an initial seeding data set (Figure 2).

**Figure 2.**
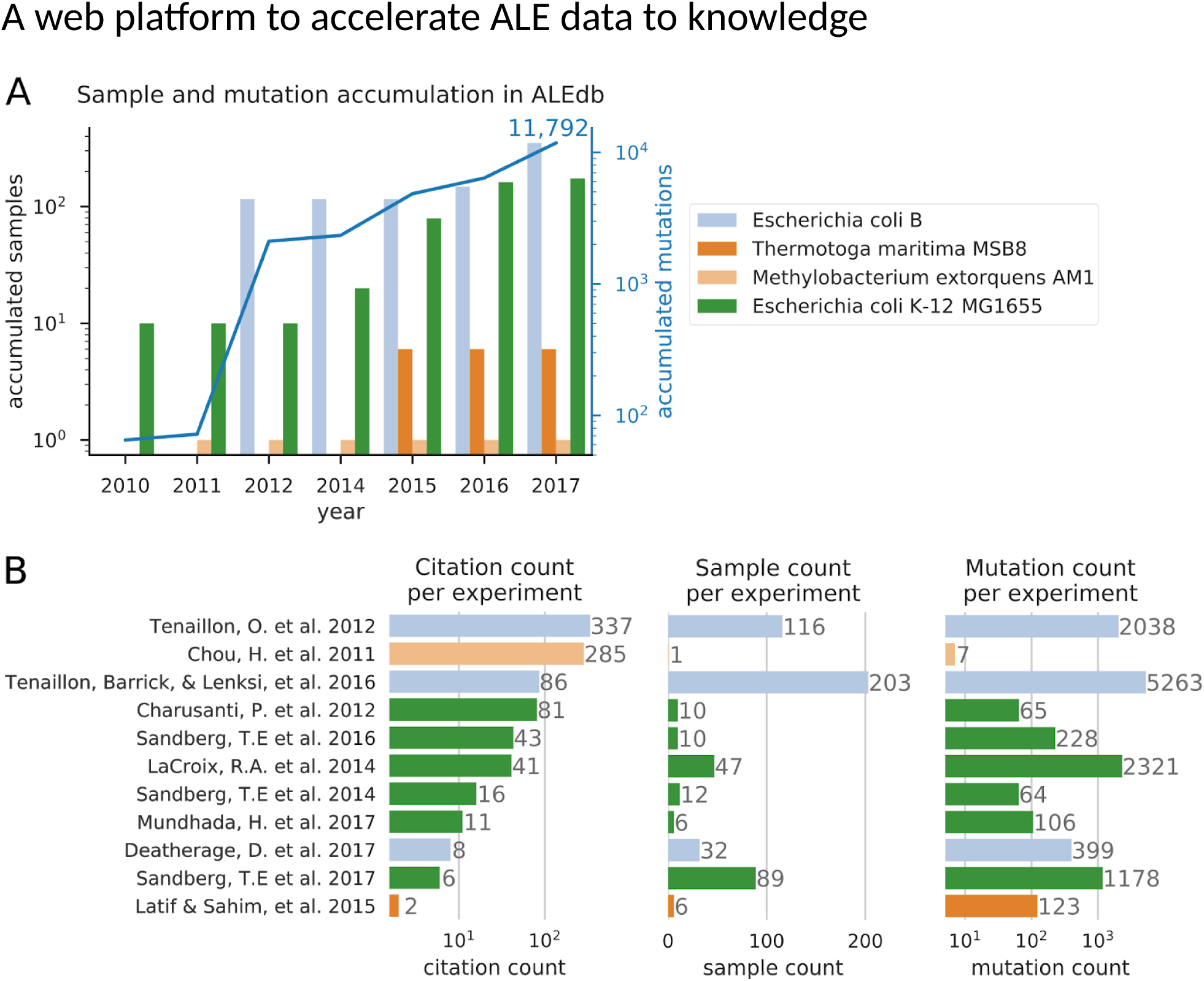
**A** A graph of the **a**ccumulation of sequenced samples and mutations in ALEdb. **B** Each publication’s sample and mutation contribution to ALEdb along with their citations at the time of ALEdb’s initial release. Citation counts were acquired from Google Scholar (scholar.google.com).

Experimental evolution studies explore the solution space of a genome optimization problem through mutational events. This element of exploration has lead to a rich diversity of published ALE experimental conditions [2]. Those experimental conditions currently represented in ALEdb are genetic perturbations [11], stress inducing environments [12], different carbon sources [13–15], and evolution duration [5]. Strains can often adapt to these conditions with a variety of different evolutionary strategies, leading to different beneficial mutations. This leads to a diversity in the mutations across ALE experiments. This rich variety of databased conditions and mutations have made ALEdb an attractive research resource, and further implementation has now made this information accessible through the web.

ALEdb’s feature set was developed in response to the challenge of accessible ALE mutation reporting for an ALE experiment pipeline [16]. ALEdb’s features enable intuitive navigation through consolidated ALE experiment data by providing two categories of features: those that describe individual ALE experiments, and those that describe all consolidated experiment data. To describe individual ALE experiments, ALEdb generates reports that detail ALE the mutation lineages, key mutations, and experimental conditions per ALE sample. To describe all consolidated ALE experiment data, ALEdb provides a mutation search feature, the ability to export the mutation data from one or more ALE experiments as spreadsheets, and an itemization of all publications that describe the databased mutations (Figure 3). ALEdb thus provides for an unmet need in the experimental evolution community: a platform to search and explore consolidated experimental evolution mutation data.

**Figure 3.**
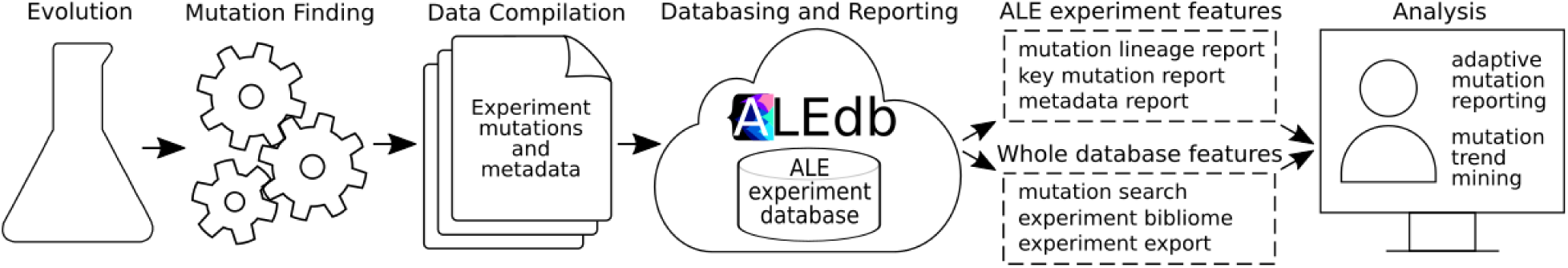
An illustration of the flow of experimental evolution data to the generation of result reports for end users and their analysis.

Mutation functional analysis is a major challenge in experimental evolution. Besides systems biology modeling methods, this task often involves searching the literature for similar results. The ALE mutations, conditions, and publications being consolidated into ALEdb can be leveraged in this work. ALEdb enables a search-driven workflow which can enhance a user’s mutation functional analysis by reporting if mutations similar to theirs have occurred in published ALE experiments. Through ALEdb’s *Search* feature, users can query for mutations of interest using multiple descriptive parameters and become aware of any databased ALE experiments that manifest similar mutations. Knowing these experiments, users can review the conditions and key mutation reports which characterize their results and refer to their associated publications through ALEdb’s *Bibliome* page. These publications ultimately describe adaptive mutations and their functional analysis, which could be leveraged by users to better understand similar mutations in their own study. ALEdb additionally includes the ability to *Export* mutation data for users interested in leveraging ALE data in applications beyond this platform (Figure 4).

**Figure 4.**
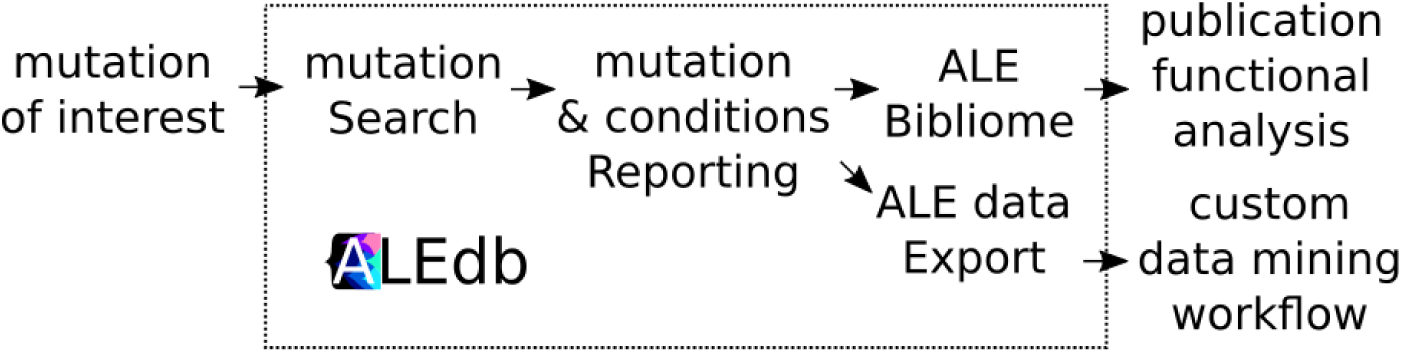
An illustration of the flow of mutation functional analysis using ALEdb. Each step within the ALEdb group is the name of a user feature on the ALEdb platform.

ALEdb’s features are described in the following sections. With ALEdb already consolidating a significant amount of ALE experiments, the final section of this study demonstrates how ALEdb can currently be used as a data resource for experimental evolution.

### Mutation search and reporting

ALEdb implements mutation *Search* to enable users to quickly find mutations of interest. Search returns a report of mutations for all databased samples according to the following mutation descriptors: gene, genome position range, mutation type, sequence change, protein change, and experiment.

Mutation search, along with most other mutation reporting mechanisms on ALEdb, present their results in the form of mutation tables (*Figure 5A*). Each ALE experiment can be described as a series of mutation sets relative to an ALE’s starting strain. Ordering sample mutation sets as columns from earliest to latest (*Figure 45, 5B*) in an ALE serves to render intuitive visualizations of temporal mutational trends. The occurrence of a mutation in a sample is annotated with an allele frequency within the intersection of the mutation row and sample column. Mutation tables therefore describe the lineage of an ALE’s final sample, or endpoint, according to the mutations that manifest during an evolution.

**Figure 5.**
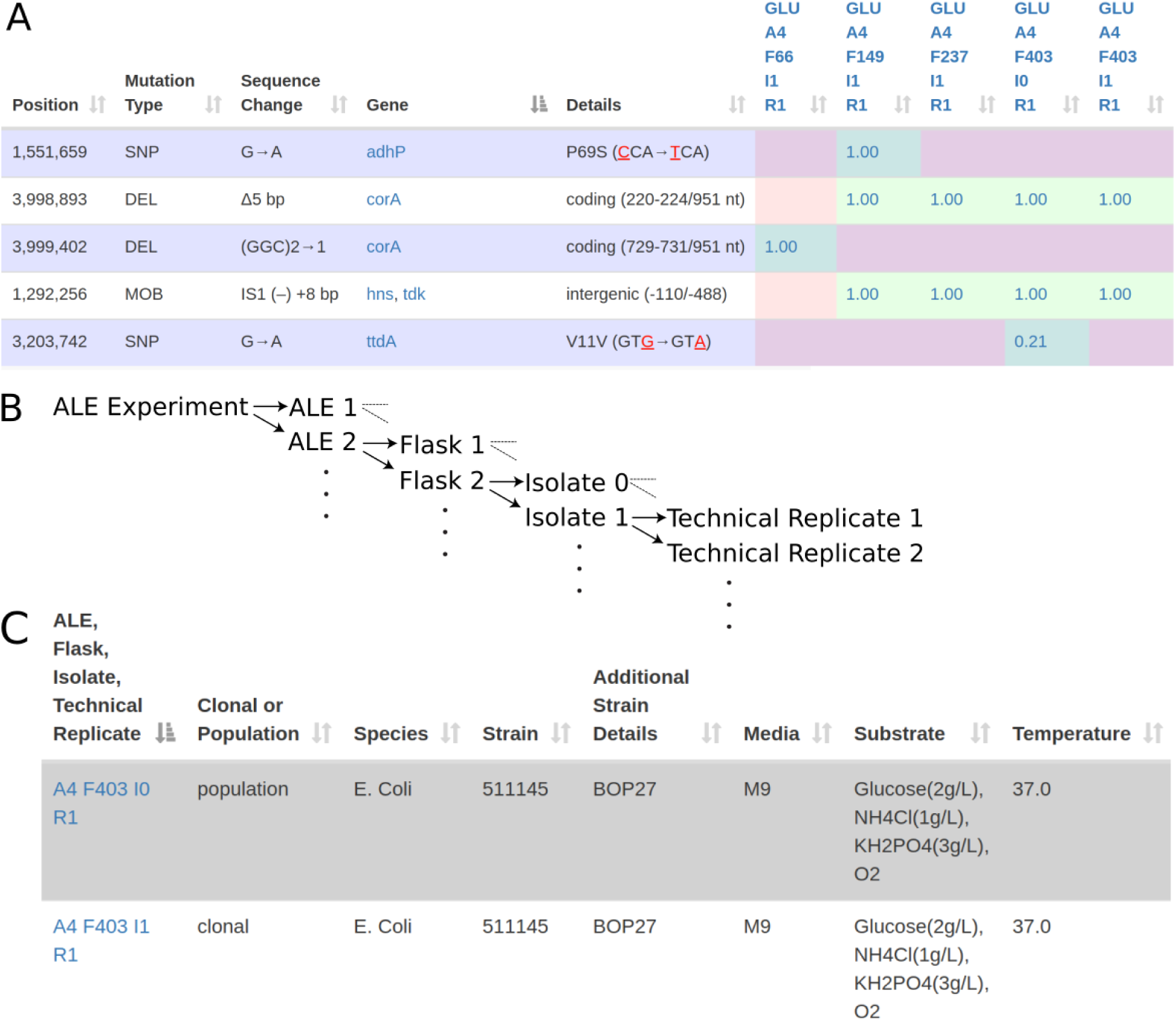
**A** An example mutation lineage report where samples are represented as columns, ordered from left to right as earliest to latest in an ALE. Rows describe the specific mutations manifested within the sample set, and values contained within cells represent the allele frequency. This format enables researchers to intuitively identify important mutational patterns, such as the fixed mutations within the *corA* gene and *hns/tdk* intergenic region. Columns are described with the experiment name, then an ALE (A#), flask (F#), isolate (I#), and technical replicate (R#) value to serialize samples. Population samples will always be described by an isolate number (I#) of 0 and will be the only sample types to carry allele frequencies less than 1.0. The information describing each mutation is generated by the mutation finding stage described in Figure 2 and details a mutations type, genomic region, and potential product effects. **B** The illustrated ordering of sample columns from left to right in the mutation lineage report. **C** An ALE experiment metadata report.

Researchers investigating ALE experiments require reporting that enables them to quickly understand which mutations are likely causal for adaptations; the mutation tables built by ALEdb are designed to meet this need. Among the many mutations that manifest within an ALE experiment, mutation rows that describe multiple alleles of a gene will cluster together according to their positions on the genome. This is illustrated with the mutated *corA* within *Figure 5A*. Due to the chronological sorting of the sample columns per ALE, a mutation that fixes across samples will manifest as an unbroken sequence of cells in a mutation row annotated with an allele frequency. This is illustrated with both the *hns/tdk* and *corA* mutations in *Figure 5A*. These two patterns are obvious to an observer and serve well to describe the adaptive mutational trends in ALE experiments.

ALE experiment mutations cannot be completely understood without considering the experiment’s conditions. ALEdb includes reports that describe an ALE experiment’s strain, substrate, and environment (*Figure 5C*). This experiment metadata can additionally be exported as spreadsheets for analysis workflows external to ALEdb.

### Consolidated ALE knowledge

A key component in the utility of ALEdb is the per experiment knowledge built from the databased mutations. The *Bibliome* feature itemizes the publications that studied the ALE mutations databased within ALEdb. Users can leverage the mutation functional analysis within these publications toward understanding any similar mutations in their experimental evolutions.

### ALE experiment mutation export

ALEdb implements an *Export* feature to give users the freedom to perform any analysis of interest on the hosted data. This feature enables users to extract one or more experiment mutation sets into comma separated value files. Users can then leverage custom analysis pipelines on the ALEdb’s data towards generating novel results.

### Automated ALE experiment key mutation reporting

ALEdb includes features that automate the reporting of established ALE adaptive mutation trends. These trends are termed *fixed* and *converged* key mutations, where each trend describes a unique pattern of mutations occurring within or across multiple ALEs in an experiment. These patterns have been used in published ALE studies to identify adaptive mutations [11–14]. The manual consolidation of adaptive mutation evidence can be prone to human error, inconsistent between researchers, and time consuming. The automation of these common analyses contributes to more consistent analysis and more accurate results.

A *fixed* mutation is one in which manifests in an ALE’s midpoint, or intermediate sample, and is propagated to all following samples in the ALE. The propagation of a mutation from their emergence to an ALE’s endpoint may describe the selection of a mutation due to its fitness benefits [13]. This analysis is only possible if an ALE experiment includes midpoint samples, providing the possibility of more than one data point per ALE mutation. The identification of *fixed* mutations is accomplished by organizing mutations according to the ALE’s sample chronology and identifying mutations that emerge in a midpoint and manifest in all following samples of the same ALE (Figure 6A). ALEdb’s *fixed* mutation reporting automates this analysis and reports results in the format described in Figure 4A.

**Figure 6.**
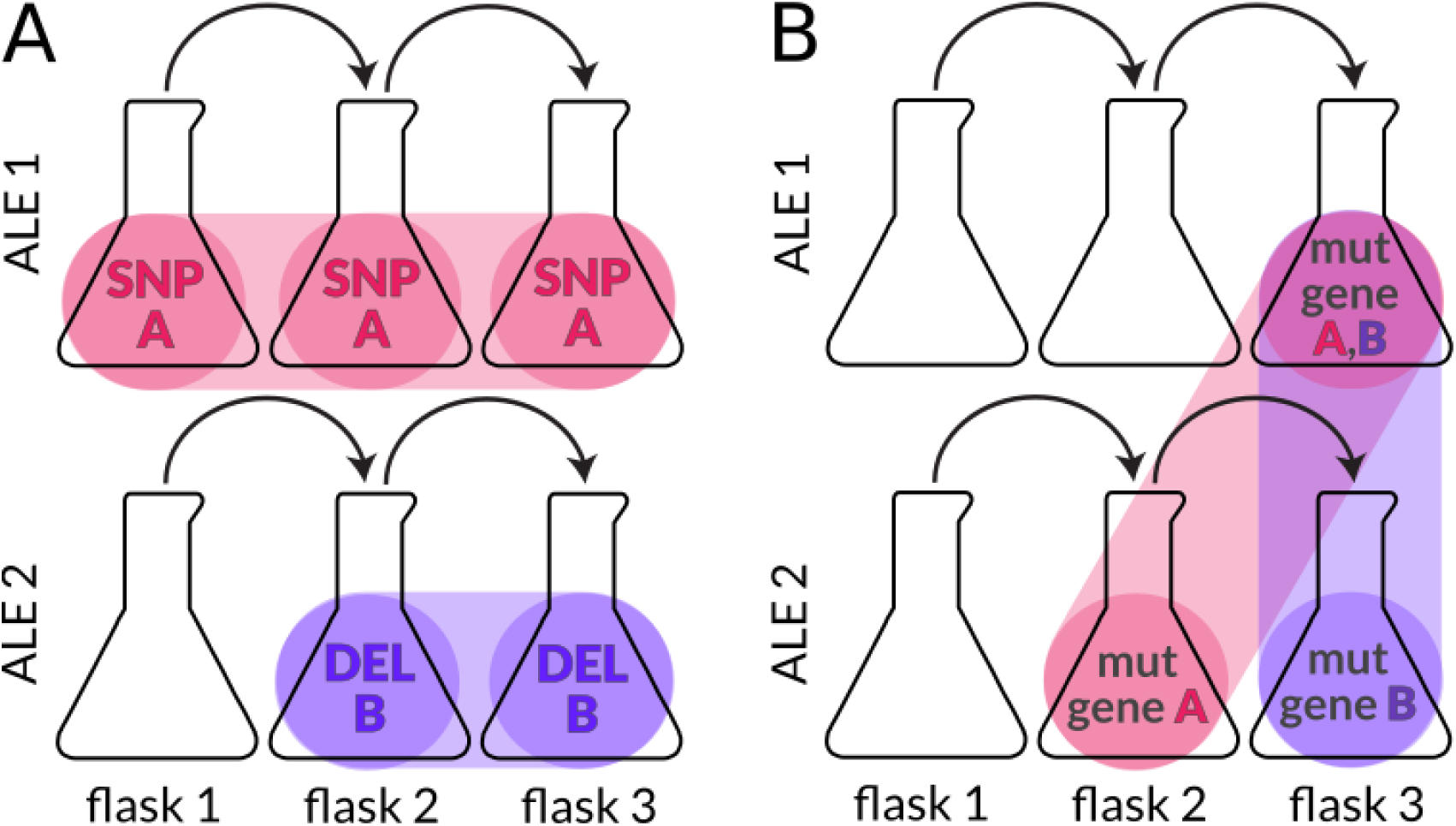
Intuition for converging and fixed mutation reports. **A** SNP A and DEL B occur in separate ALE replicates and persist through all subsequent flasks. **B** genetic targets A and B are seen to mutate across ALEs.

A *converged* mutation is one in which manifests in a genetic region seen to be mutated in multiple replicate ALEs (Figure 6B). This phenomenon describes evidence of a potential common adaptive trajectory between microbes exposed to the same conditions and has been leveraged in ALE analysis methods to more quickly identify mutations causal for adaptive phenotypes [13]. ALEdb’s *converged* mutation reporting automates this analysis and reports results in the format described in Figure 5A.

### Design and implementation

ALEdb is implemented and deployed using a standard web application technology stack and a combination of user interface technologies. ALEdb’s server-side hosts a MySQL database (https://www.mysql.com/). uses the Python based Django web framework for pre-built web application features (https://www.diangoproiect.com/). and serves the content using Gunicorn (http://gunicorn.org/) and Nginx (https://www.nginx.com/). ALEdb implements its user-interface with HTML, CSS, and JavaScript along with a combination of essential libraries, including Bootstrap (https://getbootstrap.com/). jQuery (https://iquery.com/). DataTables (https://datatables.net/), mutation-needle-plot (http://dx.doi.org/10.5281/zenodo.14561), webGL Protein Viewer (http://dx.doi.org/10.5281/zenodo.20980), and D3 (https://d3is.org/).

### Characterization and comparison of experimental evolution mutation sets

To demonstrate the potential for ALEdb as a consolidated mutation data resource for the field of experimental evolution, the experiment set databased in ALEdb and generated by the Systems Biology Research Group *(SBRG)* [5,10–15,18] was compared to experimental evolutions consolidated as two different sets. The experimental evolution mutation set from the Dettman and Kassen studies *(KEE)* [19,20] was chosen for this comparison due to its parallel nature with the that of SRBG’s. The KEE work consolidated the mutation data from 12 bacteria based experimental evolutions from the late 2000s and early 2010s, and established some of the mutation trends used in our comparisons. The Long Term Experimental Evolution *(LTEE)* is currently one of the most thoroughly studied experimental evolution mutation sets and its decades of mutation data has been released to the public [5]. The LTEE experiment set was chosen to represent a key data set for the field of experimental evolution. Mutation trends observed in previous publications were investigated across these three experiment sets [7,19,20],resulting in an overall high level of agreement across experiment sets in their recapitulation of these previously observed mutation trends.

Mutation type distributions were investigated across experiment sets to compare their endpoint proportions. The mutation finding software used by the SBRG and LTEE [21] describes mutations as single nucleotide polymorphisms (SNP), deletions (DEL), mobile element (MOB), and insertions (INS). The KEE set describes its mutations as SNPs or structural variants (SV), where SVs describe the combination of DEL, MOB, and INS mutations. SNPs are the most common mutation across all datasets, with deletions, mobile elements, and insertions being the most common structural variants across SBRG and LTEE sets respectively (Figure 7A). All experiment sets produced the same mutation type abundance order and demonstrate similar distributions (Figure 7A, Table S1).

**Figure 7.**
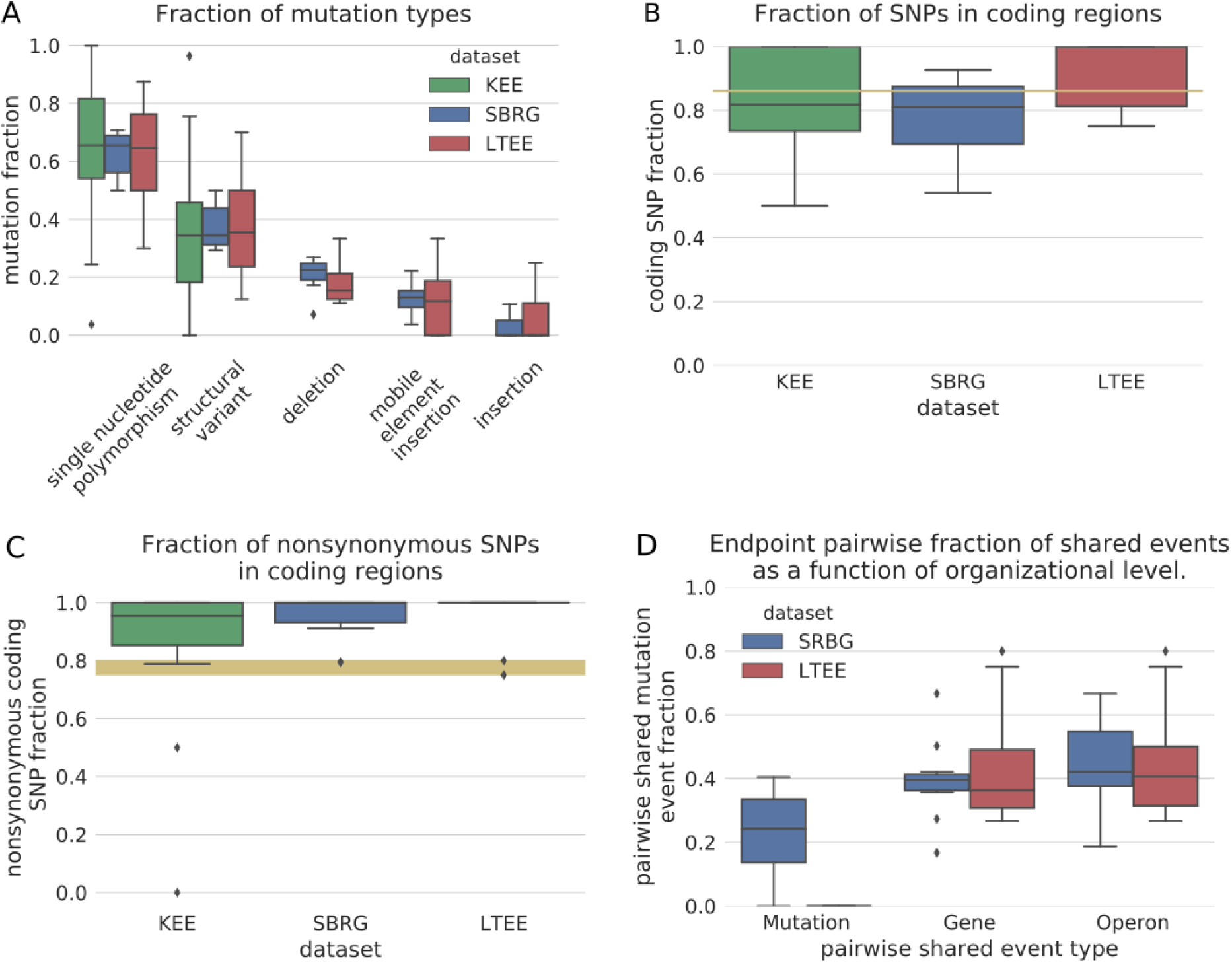
**A** Endpoint mutation type proportion distributions across experiment sets. **B** Endpoint coding SNP proportion distributions across experiment sets. **C** Endpoint synonymous SNP distributions across experiment sets. **D** Coding endpoint mutation pairwise parallelism distributions for the SBRG and LTEE experiment sets. Operons were obtained from DOOR [17].

The frequent manifestation of SNPs in experimental evolution endpoints suggests that they are highly correlated with adaptations. Previous publications investigated the selectivity of SNPs according to the density of open reading frames within bacterial genomes. They proposed that if coding SNPs were more causal for adaptations among all SNPs, their evolution endpoint proportions would be significantly larger than the proportion of coding nucleotides within the *E. coli* bacterial genome (86%) [19]. The coding SNP proportion distributions of all three experiment sets overlap the average bacterial open reading frame genome proportion and don’t provide evidence of being statistically different (Figure 7B, Table S2). The SBRG and LTEE distributions are found to be significantly different (Table S3), which may result from the different coding SNP selection characteristics in their executions.

Previous studies hypothesized that nonsynonymous SNPs are often selected as adaptive mutations in experimental evolutions [19,20]. Given a SNP, the published range for the fraction of the bacterial genome that can result in a nonsynonymous mutation is given as between 75% and 80%, depending on the species [19,20]. The LTEE and SBRG sets demonstrate significant differences from this range (Table S4). Though the KEE distribution doesn’t demonstrate this same significance, it is shown to be significantly similar to both the LTEE and SBRG sets, therefore lending evidence towards it likely being borderline significant (Table S5). Overall, the experiment sets agree in presenting evidence of selection for nonsynonymous SNPs.

A common method for finding adaptive mutations in an experimental evolution is through identifying common mutational events between replicate evolutions [7]. Similar mutations that manifest across independent replicate evolutions provide strong evidence towards a beneficial fitness effect. When the starting strain of the replicate evolutions are identical, the repeat manifestation of mutational events is known as parallel evolution [20]. The frequency of parallel evolution among replicates depends on the resolution of genomic details considered. Parallel evolution may be more likely when considering broad levels of organization, such as genes and operons, and less likely when considering higher resolutions, such as sequence positions [19]. A pairwise comparison of coding mutations between replicate evolution endpoints within experiments demonstrates that the broader the functional category of mutated genomic regions considered, the larger the fraction of shared mutation events between endpoints (Figure 7D). The KEE dataset did not include the necessary mutation details to test for parallelism between replicates and therefore could not be compared alongside the SBRG and LTEE experiments. This result recapitulate previous observations of parallelism increasing when considering higher levels of genomic organization [7,19,20] and demonstrate similar proportions of parallelism between the SBRG and LTEE datasets (Table S6).

This case study of mutation trend mining demonstrates the use of ALEdb as a source for riche experimental evolution mutation data. The results demonstrate the agreement of the consolidated experimental evolution mutation data with previously established trends and the agreement of trends across data generated from different groups.

## CONCLUSION

ALEdb works to serve the current need for a mutation database in the field of experimental evolution. It is a platform designed for the integration and reporting of ALE mutation datasets and currently integrates the mutation data and published materials of eleven published ALE experiments. Additionally, multiple features are implemented within ALEdb to enable intuitive navigation and analysis. Finally, the case study included in this work demonstrates the potential for ALEdb as a mutation data resource for investigating experimental evolution trends.

ALEdb will continue to be developed to meet the needs of consolidating, reporting, and navigating ALE experiment data. This initial release of ALEdb considers previously generated mutation datasets. ALEdb will continue to grow with future inclusion of published ALE experiment results from currently contributing and new research organizations.

## AVAILABILITY

ALEdb is freely available online at http://aledb.org and can be accessed with a JavaScript-enabled browser.

## METHODS

### Mutation finding pipeline

Mutation data currently hosted on ALEdb are generated by the *breseq* mutation finding pipeline [21]. Being that these samples come from different projects, various version of breseq were used in their mutation data generation. The sequencing reads used to generate the mutation data were subjected to quality control through either *FastQC* and *FastX-toolkit* or *AfterQC* [22–24].

### Experimental evolution sample selection for case study

In comparing the endpoints of the SBRG, KEE, and LTEE experiment sets, strategies for normalizing between experiments of different replicate evolution counts and lengths were necessary. To normalize for approximate evolution length, LTEE samples at 2000 generations were considered endpoints. To normalize for different replicate evolution counts per experiment within the SRBG experiment sets, the average of each result across replicates is taken to represent an experiment. Additionally, no samples containing hypermutator strains were included. Though hypermutator strains are informative, they represent a cellular state that isn’t easily comparable to strains with their DNA repair mechanisms intact.

## ACKNOWLEDGEMENTS

This work was supported by the Novo Nordisk Foundation Grant Number NNF NNF10CC1016517.

## CONTRIBUTIONS

PVP, BOP, and AMF designed the study. PVP and DG consolidated the data and implemented the software. PVP, BOP, and AMF wrote the paper.

